# The mitochondrial matrix protein cyclophilin D contributes to deficits in parvalbumin interneurons in schizophrenia

**DOI:** 10.1101/2023.05.19.541499

**Authors:** John T. O’Brien, Sophia P. Jalilvand, Neha A. Suji, Aarron Phensy, Rohan K. Jupelly, Juliet M. Mwirigi, Hajira Elahi, Theodore J. Price, Sven Kroener

**Author notes:** Correspondence should be addressed to SK.

## Abstract

**BACKGROUND:** Cognitive deficits in schizophrenia are linked to dysfunctions of the dorsolateral prefrontal cortex (DLPFC), including alterations in parvalbumin (PV)-expressing interneurons (PVIs). Redox dysregulation and oxidative stress may represent convergence points in the pathology of schizophrenia, causing dysfunction of GABAergic interneurons and loss of PV. Here, we show that the mitochondrial matrix protein cyclophilin-D (CypD), a critical initiator of the mitochondrial permeability transition pore (mPTP) and modulator of the intracellular redox state, is altered in PVIs in schizophrenia.

**METHODS:** Western blotting was used to measure CypD protein levels in postmortem DLPFC specimens of schizophrenic (n=27) and matched comparison subjects with no known history of psychiatric or neurological disorders (n=26). In a subset of this cohort, multilabel immunofluorescent confocal microscopy with unbiased stereological sampling methods were used to quantify 1) numbers of PVI across the cortical mantle (20 control, 15 schizophrenia) and 2) PV and CypD protein levels from PVIs in the cortical layers 2-4 (23 control, 19 schizophrenia).

**RESULTS:** In schizophrenic subjects, the overall number of PVIs in the DLPFC was not significantly altered, but in individual PVIs of layers 2-4 PV protein levels decreased along a superficial-to-deep gradient when compared to unaffected comparison subjects. These laminar-specific PVI alterations were reciprocally linked to significant CypD elevations both in PVIs and within total DLPFC gray matter.

**CONCLUSIONS:** Our findings support previously reported PVI anomalies in schizophrenia and suggest that CypD-mediated mPTP formation could be a potential contributor to PVI dysfunction in schizophrenia.

## INTRODUCTION

Disruptions of cognitive processes, such as impairments in working memory, are important symptoms in the pathophysiology of schizophrenia which often present prior to the onset of psychosis [1-3]. Converging lines of evidence link these disturbances to dysfunctional microcircuits in the dorsolateral prefrontal cortex (DLPFC), where reciprocal connections between pyramidal neurons and parvalbumin (PV)-expressing interneurons (PVIs) are altered [4, 5]. Abnormalities in PVIs, including a reduced density of immunoreactive neurons as well as lower PV mRNA and protein levels, constitute one of the most robustly replicated findings in human postmortem studies of schizophrenia [6, 7]. Although the exact cause of these changes is unclear, redox dysregulation and oxidative stress may present a convergence point for pathologies in schizophrenia, causing dysfunction of GABAergic interneurons and loss of PV [8, 9]. As the primary source of reactive oxygen species (ROS) in neurons, mitochondria are both crucial contributors and amplifiers of intracellular oxidative stress in pathological states [10-13]. Transcriptomic, proteomic, and metabolomic studies have shown that schizophrenia is associated with changes in mitochondrial genes, proteins, and pathways involved in energy production, ROS formation, and antioxidant defense systems [14-17]. Cell type-specific studies have further linked these mitochondrial alterations to deep layer 3 pyramidal neurons and PVIs in the DLPFC [18, 19]. Fast-spiking PVIs are particularly vulnerable to oxidative stress-driven impairment due to their enriched mitochondrial profile necessary to satisfy their high energy demands [11]. During times of high oxidative stress, mitochondria transition to a more permeable state, leading to the translocation of the mitochondrial matrix protein cyclophilin D (CypD) to the inner mitochondrial membrane [20-22]. This translocation acts as a key trigger for the opening of the mitochondrial permeability transition pore (mPTP), a large, nonselective channel that plays a critical role in regulating mitochondrial function and cellular homeostasis [23-25]. Under conditions of chronic stress, elevated levels of CypD can lead to maladaptive changes in mitochondria, including decreased adenosine triphosphate (ATP) production and increased release of ROS into the cytosol [22, 25, 26]. These alterations contribute to a feed-forward cycle of ROS generation and release, metabolic changes, and eventual loss of cell function [23, 24]. Despite mitochondria’s importance in redox regulation and extensive evidence for the contribution of oxidative stress to the pathophysiology of schizophrenia [8, 9, 12, 27], the role of CypD in these processes remains largely unclear [28]. We hypothesized that PVI alterations in schizophrenia may be associated with increased CypD protein levels. To test this idea, we investigated the relationship between CypD and PVIs in human DLPFC tissue collected from postmortem schizophrenic patients and unaffected comparison subjects. Using a combined approach of western blotting, immunohistochemistry, and confocal fluorescence microscopy with unbiased stereological sampling methods, we measured CypD expression levels in DLPFC gray matter homogenates, and quantified PV and CypD protein levels in PVIs from cortical layers 2-4. Our findings suggest that schizophrenia is linked to changes in DLPFC mitochondria which may be part of a larger network of pathologic cellular processes tying oxidative stress to alterations in cortical PVIs.

## METHODS AND MATERIALS

### Human Subjects

Human tissue samples were sourced from the National Institute of Health (NIH) NeuroBioBank (NBB), with contributions from the following biorepositories: Harvard Brain Tissue Resource Center, University of Maryland Brain and Tissue Bank, University of Miami Brain Endowment Bank, Sepulveda Research Corporation, and the Mount Sinai/JJ Peters VA Medical Center. Brain specimens of area 9 and 46 (n=108) were obtained as frozen (n=54), formalin-fixed (n=46), and formalin-fixed paraffin-embedded (FFPE, n=8) tissue preparations. For 42 subjects (22 control, 20 schizophrenia), we received both frozen and formalin-fixed preparations. For 8 subjects (4 control, 4 schizophrenia), we received both frozen and FFPE preparations. For 4 subjects (1 control, 3 schizophrenia), we received only frozen tissue. For 4 subjects (1 control, 3 schizophrenia), we received only formalin-fixed tissue. Subjects with schizophrenia were matched as closely as possible for age, sex, and postmortem interval (PMI) to unaffected comparison subjects, and tissue samples from members of a pair/triad were always processed together. Subject groups did not differ in mean age or PMI. Brain specimens were allocated to western blot (n=53) and immunohistochemistry (n=42) experiments as outlined in Table 1. From the 46 formalin-fixed samples (23 control, 23 schizophrenia) and 8 FFPE samples (8 control, 8 schizophrenia) available for immunohistochemistry experiments, 35 formalin-fixed samples (20 control, 15 schizophrenia), and 7 FFPE samples (4 control, 3 schizophrenia) were used in the final reported analysis. The other 11 formalin-fixed samples (3 control, 8 schizophrenia) and 1 FFPE sample (control) were excluded because their number of immunolabeled PV+ somata was more than 4 standard deviations below their respective group average (Supplemental Figure 1). From the 54 frozen samples (27 control, 27 schizophrenia) available for western blot experiments, 1 sample (control) was excluded from the analysis because CypD and GAPDH protein intensity measurements both were more than 4 standard deviations below the group mean.

**Table 1:** Summary of Human Subject Characteristics

| Characteristic | <u>Study and Group</u> |  |  |  |  |  |  |  |
| --- | --- | --- | --- | --- | --- | --- | --- | --- |
|  | <u>Western Blotting</u> |  |  |  | <u>Immunofluorescence</u> |  |  |  |
|  | Comparison Group<br>(N=26) |  | Schizophrenia Group<br>(N=27) |  | Comparison Group<br>(N=23) |  | Schizophrenia Group<br>(N=19) |  |
|  | N | % | N | % | N | % | N | % |
| Sex |  |  |  |  |  |  |  |  |
| Male | 20 | 76.9 | 21 | 77.8 | 19 | 82.6 | 15 | 78.9 |
| Female | 6 | 23.1 | 6 | 22.2 | 4 | 17.4 | 4 | 21.1 |
| Tissue Preparation |  |  |  |  |  |  |  |  |
| Frozen | 26 | 100 | 27 | 100 | 0 | 0 | 0 | 0 |
| Formalin-Fixed | 0 | 0 | 0 | 0 | 20 | 87.1 | 15 | 78.9 |
| FFPE | 0 | 0 | 0 | 0 | 3 | 12.9 | 4 | 21.1 |
|  | Mean | SD | Mean | SD | Mean | SD | Mean | SD |
| Age | 50 | 8.4 | 47.9 | 10.4 | 49.7 | 8.7 | 50.6 | 8.7 |
| PMI | 20.5 | 6.1 | 20.4 | 10.9 | 19.8 | 5.4 | 20.1 | 10.1 |

Abbreviations: FFPE, formalin-fixed paraffin-embedded; PMI, post-mortem interval. Values are mean +/- SD; $p > 0.5$ for all comparisons.

**Figure 1:**
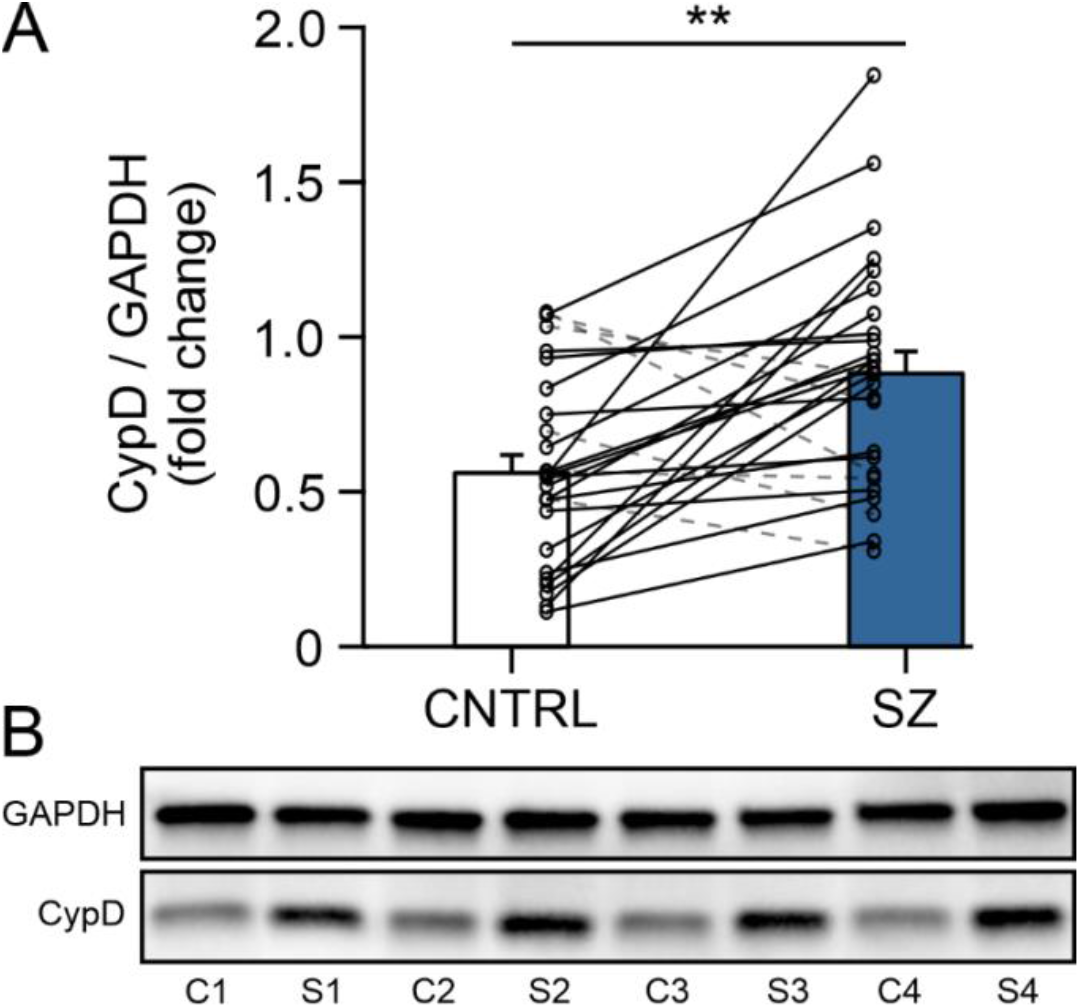
CypD protein levels are elevated in postmortem tissue of the DLPFC from subjects with schizophrenia. A) Hybrid bar graph and scatter plot showing that in 21/26 matched pairs subjects with schizophrenia (SZ) showed significantly higher GAPDH-normalized CypD protein levels than unaffected comparison subjects (CNTRL). Error bars = SEM, p < 0.01. B) Lanes from a representative Western blot loaded with tissue from 4 matched comparison (C1-4) and schizophrenia (S1-4) subject pairs. Abbreviations: CypD, cyclophilin D; DLPFC, dorsolateral prefrontal cortex; GAPDH, glyceraldehyde-3-phosophate dehydrogenase

### Tissue Processing & Immunohistochemistry

FFPE and formalin-fixed DLPFC tissue samples from 42 matched subjects (15 pairs + 4 triads) were sectioned rostral-caudally at either 40 µm (formalin-fixed) or 15 µm (FFPE) on a sliding microtome. Because of the difference in thickness only formalin-fixed sections were used in experiments that compared densities of PVI across the layers. Three sections per subject were selected for immunohistological staining, with each section being separated by ∼400 µm to ensure a representative sampling of the region of interest and to minimize experimental variance. Tissue from subject pairs/triads were processed together using antibodies against PV, CypD, and counterstained with Neuronal Nuclear marker protein (NeuN) or 4′,6-diamidino-2-phenylindole (DAPI). See Supplemental Methods for details.

### Microscopy

Data were collected on an Olympus IX83 motorized inverted microscope equipped with a FV3000RS confocal laser scanning module, GaAsP detector photomultiplier tube, ultrasonic automated stage with XYZ linear encoders, and using 1.25X (numerical aperture, N.A. 0.04), 20X (N.A. 0.75), and 100X (N.A.1.35) air or oil immersion objectives. Z-stacked images were collected at low magnification (20X air objective, 1024x1024 frame size, 30 µm thickness, 2 µm step size) and high magnification (100X silicone objective, 1024x1024 frame size, 0.25 µm step size) using FV3000 Fluoview Acquisition and Analysis software. Exposure times during image capture were optimized to prevent saturated conditions for each channel within each image stack and differences in exposure times were corrected before processing.

### Stereological Methods and Image Capture

To map cellular and subcellular changes, we integrated a stratified stereologic sampling scheme informed by a previously published protocol [29] which provided a systematic, unbiased approach in order to 1) chart the distribution of PVIs across cortical layers and 2) randomly select individual PVIs in specific layers for fluorescence intensity measurements of PV and CypD. The boundaries of the six cortical layers were estimated based on the distance from the pial surface to the white matter based on measurements made in NeuN-stained sections, following definitions based on [30, 31]: 1 (pia-10%), 2 (10%–20%), 3 (20%–50%), 4 (50%–60%), 5 (60%–80%) and 6 (80%-gray/white matter border). For all subjects, a minimum of 5 PVI from layer 2, 15 PVI from layer 3, and 10 PVI from layer 4 were selected across three coronal sections, totaling ∼30 PVI per subject. See Supplemental Methods for details.

### Image Processing and Analysis

Images were imported into Imaris Software (Oxford Instruments, version 9.0.2) and pixels from previously acquired z-stacks were converted into 3D voxels. This information was then used to reconstruct the original 3D object that spanned across the z-stacks. Images were then subjected to intensity histograms for each channel and independently adjusted to identical settings across all subjects allowing PV+ and CypD+ cells to be masked under identical conditions. The surface and spot detection functions were used to count PVI across all six layers of the neocortex. For each subject, the number of PVI was then averaged for each layer within each coronal section. For individual PVIs, a region of interest was traced across the depth of the z-plane and the surface wizard was used to mask individual channels. Mean fluorescence intensity measurements were then recorded for PV and CypD and values were averaged for each subject.

### Western blotting

Frozen and frozen-pulverized DLPFC tissue samples from 53 matched subjects (25 pairs + 1 triad) were processed using antibodies against CypD and glyceraldehyde-3-phosphate dehydrogenase (GAPDH), then visualized with chemiluminescent substrate. CypD band strength was normalized to GAPDH protein intensity. All samples were run at least twice, and values were averaged for statistical analysis. See Supplemental Methods for details.

### Statistics

Analysis of covariance (ANCOVA) models were used to compare the dependent variables between schizophrenia and unaffected comparison subject groups. Age, sex, PMI, and tissue preparation were included as covariates. Nonsignificant covariates were excluded in the final reported analyses. For western blot experiments, a one-way between-subjects ANCOVA was performed to evaluate CypD protein expression levels in the DLPFC. For immunohistochemistry experiments, three mixed-design (split-plot) ANCOVAs were performed to evaluate the number of PVI across the cortical mantle, or PV and CypD intensity measurements from individual neurons, respectively. Clinical diagnosis was used as the between-subjects factor with repeated-measures on comparisons of cortical layers. If Mauchly’s test indicated that the assumption of sphericity had been violated (p < 0.05), then degrees of freedom were corrected using either Greenhouse-Geisser (ε < 0.75) or Huynh-Feldt (ε > 0.75) estimates of sphericity. Diagnosis and diagnosis-by-layer effects were tested using *F* tests, followed by Bonferroni adjusted *p* values in each layer.

## RESULTS

### CypD expression levels are elevated in the DLPFC for subjects with schizophrenia as measured by western blot

We used frozen tissue samples from 53 subjects (26 control, 27 schizophrenia) to measure CypD protein expression levels in the DLPFC using western blot (Figure 1). Expression levels were normalized to the housekeeping protein GAPDH to account for potential differences in protein loading. Mean values for GAPDH did not significantly differ (F_1,48_ = 1.050, p = 0.311, η_p_^2^ = 0.021) between tissue from control subjects (1.11E+7 +/- 7.91E+6) and subjects with schizophrenia (9.27E+6 +/- 6.10E+6; Figure 1). There was a significant main effect of diagnosis on GAPDH-normalized CypD expression levels (F_1,48_ = 10.568, p = 0.002, η_p_^2^ = 0.180), with a 34.9% increase in the CypD/GAPDH ratio for schizophrenia subjects (0.847, +/- 0.058) compared to unaffected comparison subjects (0.569, +/- 0.059; Figure 1). Taken together, these elevations in CypD protein levels provide evidence for an altered mitochondrial redox state in the DLPFC of subjects with schizophrenia.

### The laminar distribution of PVIs across the DLPFC does not differ between schizophrenic and matched comparison subjects

To determine if cortex-wide increases in CypD are associated with PVI abnormalities in subjects with schizophrenia, we mapped the distribution of immunoreactive PV+ somata across all six layers of the DLPFC in 35 matched comparison subjects (20 control, 15 schizophrenia) and quantified the number of PVI somata in each cortical layer (Figure 2). There was a significant main effect for the cortical layer variable on the number of PVI somata (F_1.93,57.98_ = 12.137, p < 0.001, η_p_^2^ = 0.288), indicating that the distribution of PVIs significantly varied across cortical layers (Figure 3). In both schizophrenia and matched comparison subject groups, the distribution of PVIs across the cortical layers were as follows: ∼ 9% in layer 2, 38% in layer 3, 31% in layer 4, 16% in layer 5, and 5% in layer 6. We detected little to no PV labeling in cortical layer 1 (Figure 3). This distribution pattern is consistent with previous reports about the laminar distribution of PVIs in human and monkey PFC [30, 32]. We did not find a significant main effect of diagnosis on the number of PVI somata (F_1,30_ = 2.399, p > 0.131, η_p_^2^ = 0.074), nor was there a significant interaction between cortical layer and diagnosis (F_1.93,57.98_ = 2.163, p > 0.151, η_p_^2^ = 0.06), indicating that on average the number of PVI did not significantly differ between tissue from subjects with schizophrenia and unaffected control subjects (Figure 3).

**Figure 2:**
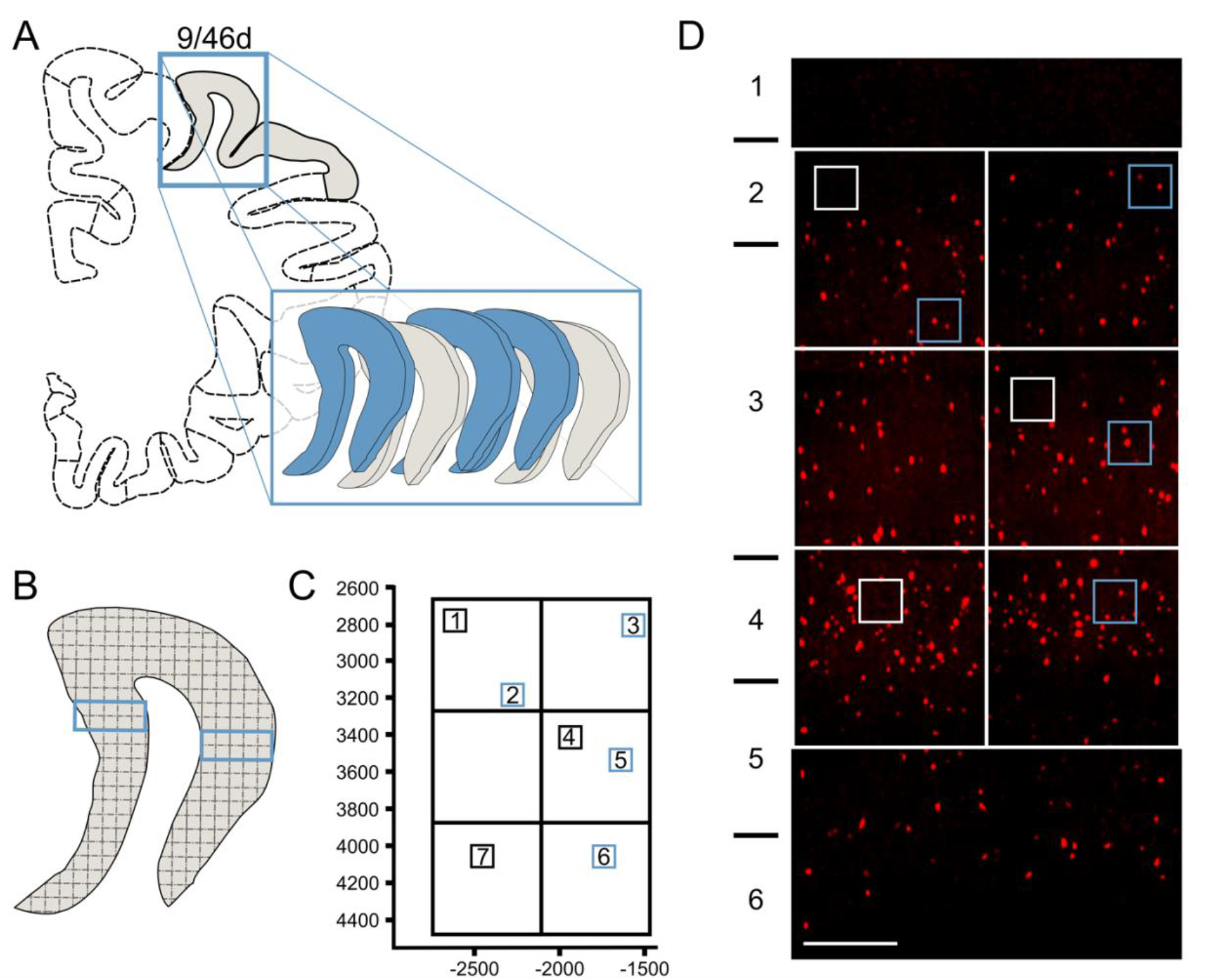
Stereological methods for systematic random sampling of PVIs in the human DLPFC. A) Coronal section at level of DLPFC with Brodmann’s areas 9/46. Inset is area 9 which was sectioned coronally on a freezing microtome and 3 sections per subject (blue) were chosen for antibody staining. B) Sections were imaged on a confocal microscope using a 1.25X objective and 636x636 µm random sampling grids were overlayed across the gray matter. Two regions of the gray matter (inset blue boxes) were chosen pseudorandomly to cover comparable regions in areas 9/46 in each subject, and confocal z-stacks were acquired to cover the extent from the pial surface to the white matter border. C) A MATLAB script was used to generate random grid coordinates for selection of individual PVI in layers 2-4 within a 125x125 sampling grid (inset boxes). D) Confocal z-stacked and spliced image using a 20X showing distribution of PVI (red) across the cortical mantle. Grid coordinates which included PV-positive somata (blue insets) were accepted for further analysis of PVI and cyclophilin-D at higher magnification, whereas grids that did not include PVI (white insets) were not imaged further. Abbreviations: PVI, parvalbumin interneuron; DLPFC, dorsolateral prefrontal cortex; PV, parvalbumin; CypD, cyclophilin D

**Figure 3:**
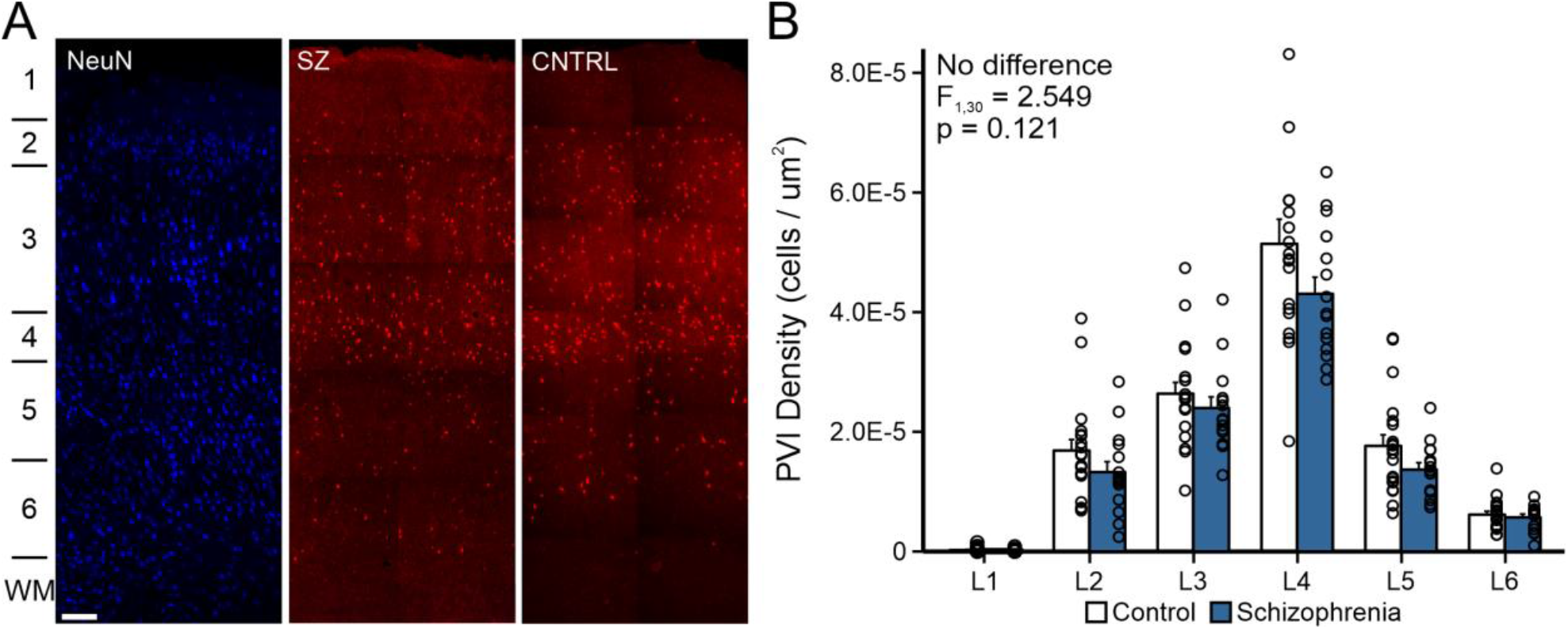
The laminar distribution of PVIs is not altered in schizophrenia. A) Representative z-stack projections (using a x 20 objective) showing the laminar organization of the NeuN (left) and PVI in postmortem tissue from unaffected comparison subjects (middle, CNTRL) and subjects with schizophrenia (right, SZ). The approximate location of cortical layers is indicated on the left. Scale bar = 200 um. B) Bar graph showing the distribution of PVIs per square micron in each cortical layer for unaffected control subjects (white bars) and subjects with schizophrenia (grey bars). Error bars = SEM. Abbreviations: PVI, parvalbumin interneuron; NeuN, neuronal marker

### PVI show a decreasing gradient of PV expression across layers 2-4 that is inversely related to an increasing gradient of CypD expression

We used tissue from the same 35 subjects as in Experiment 3.1 (20 control, 15 schizophrenia), plus an additional 7 FFPE samples (3 control; 4 schizophrenia) to quantify expression of PV and CypD at the soma of 1,266 randomly sampled PVIs (690 control, 576 schizophrenia, ∼30 PVI per subject) from layers 2-4. Our results revealed a significant main effect of diagnosis on somatic PV intensity (F_1,36_ = 103.637, p < 0.001, η_p_^2^ = 0.742) (Figure 4). In addition, although there was no main effect of cortical layer on somatic PV expression (F_1.957,70.436_ = 1.759, p = 0.180, η_p_^2^ = 0.047), there was a significant interaction between diagnosis and cortical layer (F_1.95,70.43_ = 6.203, p < 0.01, η_p_^2^ = 0.147). Post hoc analyses using Bonferroni-corrected t-tests showed that in subjects with schizophrenia PV levels were significantly decreased in PVI from layer 2 (-18.56%, F_1,36_ = 35.875, p < 0.001, η_p_^2^ = 0.499, Figure 4A, B), layer 3 (-15.41%, F_1,36_ = 38.701, p < 0.001, η_p_^2^ = 0.518, Figure 4C, D), and layer 4 (-25.3%, F_1,36_ = 109.807, p < 0.001, η_p_^2^ = 0.753, Figure 4E, F). Our results also revealed a significant main effect of diagnosis on CypD intensity (F_1,36_ = 60.471, p < 0.001, η_p_^2^ = 0.627) (Figure 4). Although there was no main effect of cortical layer on somatic CypD expression (F_1.93,69.70_ = 1.046, p > 0.354, η_p_^2^ = 0.028), there was a significant interaction between diagnosis and cortical layer (F_1.93,69.70_ = 5.124, p < 0.01, η_p_^2^ = 0.125). Post hoc analyses showed that in tissue from subjects with schizophrenia CypD protein levels in the somata of PVI were significantly increased in layer 2 (+10.08%, F_1,36_ = 14.322, p < 0.01, η_p_^2^ = 0.285, Figure 4A, B), layer 3 (+13.04%, F_1,36_ = 45.803, p < 0.001, ηp^2^ = 0.560, Figure 4C, D), and layer 4 (+19.34%, F_1,36_ = 61.60, p < 0.001, η_p_^2^ = 0.629, Figure 4E,F). Taken together our data indicate that PVIs in tissue from subjects with schizophrenia show laminar-specific reductions in PV that inversely correlated with rising CypD protein levels, suggesting that PVIs in layers 2-4 of the DLPFC are particularly vulnerable to mitochondrial pathology.

**Figure 4:**
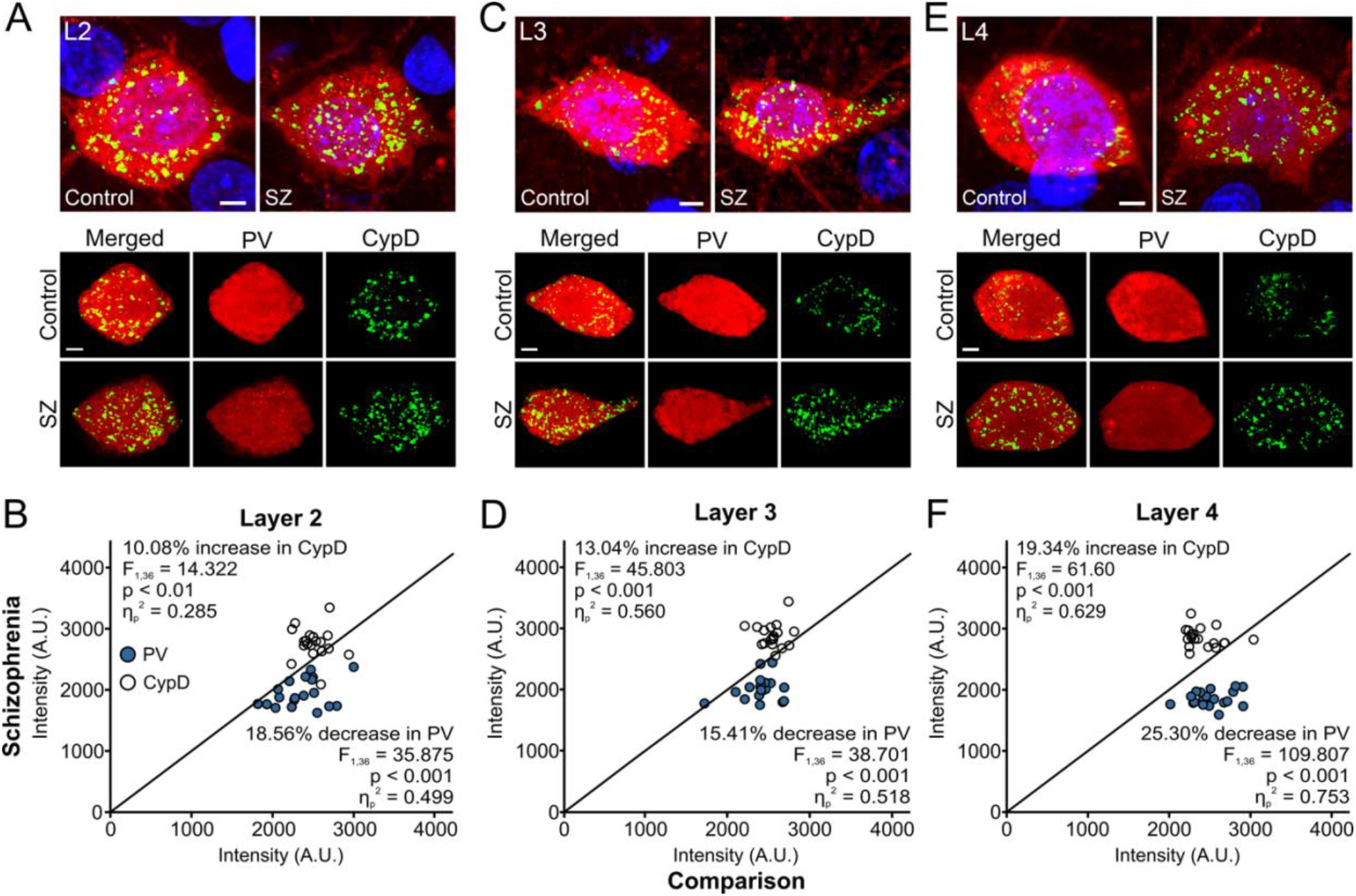
Colocalization of PV and CypD in the somata of cortical PVIs from postmortem tissue of unaffected subjects and subjects with schizophrenia. A, C, E) Representative z-stacked confocal images (using a x 100 objective) of DLPFC tissue sections immunolabeled for PV (red) and CypD (green), counterstained with DAPI (blue), in layers II (L2; A), III (L3; C), and IV (L4; E). B, D, F) Reductions in PV levels parallel increased levels of CypD at the somata of PVIs in layers 2-4 of subjects with schizophrenia. Unity plots showing the relationship between PV and CypD expression in PVIs from cortical layer 2 (B), layer 3 (D) and layer 4 (F). Each data point reflects a subject pair/triad (15 pairs + 4 triads total). Data points below the unity line indicate lower PV values and data points above the line indicate higher CypD values in the schizophrenia subject relative to the matched comparison subject. The effect size (partial eta squared, ηp2), determines the proportion of variance in the dependent variable (PV or CypD intensity) that is explained by the between-subjects independent variable (clinical diagnosis). In schizophrenic subjects, there was a superficial-deep laminar gradient of declining PV levels in layers 2-4, which was accompanied by a corresponding gradient of increased CypD levels in the same neurons. Abbreviations: PV, parvalbumin; CypD, cyclophilin D; PVI, parvalbumin interneuron; DLPFC, dorsolateral prefrontal cortex; DAPI, 4′,6-diamidino-2-phenylindole

## DISCUSSION

In this study, we investigated the relationship between the expression of the mitochondrial matrix protein CypD and alterations in PVIs in postmortem tissue of the DLPFC from schizophrenic and unaffected comparison subjects. In the DLPFC of subjects with schizophrenia, we observed elevated CypD levels in gray matter homogenates. These changes were accompanied by reduced PV expression in PVIs along a superficial-to-deep gradient in layers 2-4, as well as inversely related elevations in CypD levels in the same neurons, without significant alterations in the total number of PV+ cells.

The PVI anomalies in subjects with schizophrenia that we describe here are consistent with a large body of literature that points to alterations in cortical PVIs as a hallmark in the pathology of the disease [6, 7, 33, 34]. Several previous studies reported reduced numbers of PVIs in the prefrontal cortex [35-38] and other brain regions including the hippocampus [39, 40], entorhinal cortex [41], and thalamus [42]. However, more recent studies suggest that the numbers of PVIs in the DLPFC remain unchanged in schizophrenia; instead, alterations occur at the level of gene expression and protein synthesis, which can contribute to lower PV expression and cause cells to fall below a detection threshold [5, 33, 43-47]. Our immunohistochemical analyses are consistent with this interpretation, and also indicate an important role for mitochondrial dysfunction in the alterations in PVI in schizophrenia (Figure 5).

**Figure 5:**
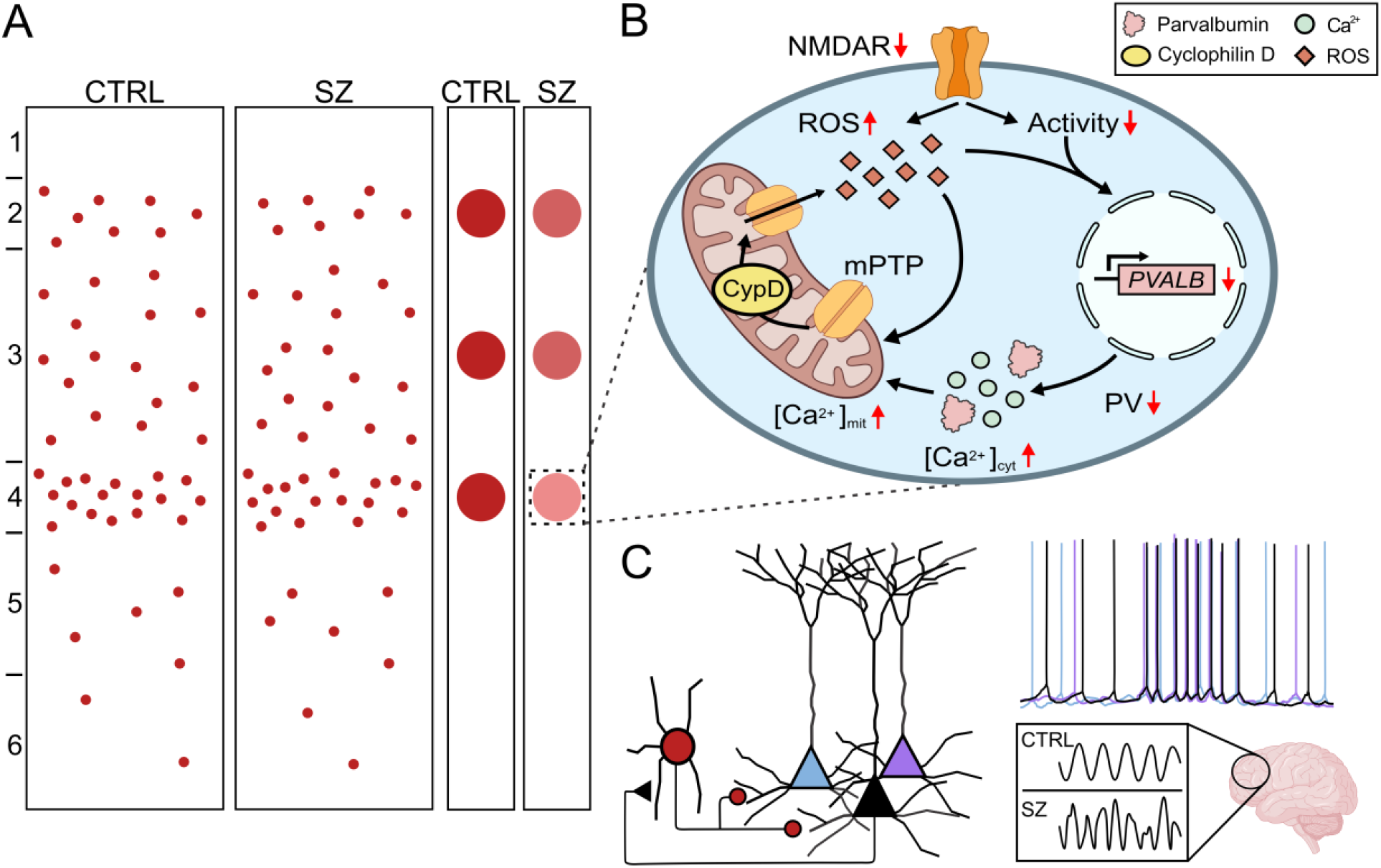
Summary of findings and association between CypD and PV and their potential role in alterations in PVI and cognitive dysfunction in schizophrenia. A) The density of PVIs in the DLPFC is not altered in schizophrenia (red dots in left columns), but PVIs in layers 2-4 exhibit a superficial-to-deep decline in PV protein levels (red dots in right columns) which are reciprocally linked to elevations in CypD. B) Patients with schizophrenia show mitochondrial impairments and heightened oxidative stress. Oxidative and other cellular stressors (including increased levels of cytosolic calcium, [Ca]cyt) promote the translocation of CypD to the inner mitochondrial membrane and facilitate opening of the mPTP. CypD-mediated mPTP opening increases production of ROS and decreases ATP synthesis, causing disruptions in cellular energy metabolism, calcium homeostasis, and antioxidant defense mechanisms. These changes can directly or indirectly affect PV expression and function, further affecting a cell’s ability to buffer cytosolic calcium. C) Deficits in supragranular DLPFC PVI may affect the synchrony of gamma oscillations that support cognitive functions. Disruptions in DLPFC gamma activity are predictor for the functional outcomes in patients with schizophrenia. Abbreviations: CypD, cyclophilin D; PV, parvalbumin; PVI, parvalbumin interneuron; DLPFC, dorsolateral prefrontal cortex; mPTP, mitochondrial permeability transition pore; ROS, reactive oxygen species, ATP, adenosine triphosphate

Multiple lines of evidence indicate that oxidative stress and redox dysregulation may be critical factors in schizophrenia’s pathophysiology across all phases of the illness [17-19, 48-54]. For example, a recent in vivo study using magnetic resonance spectroscopy found significant reductions in the ratio of oxidized to reduced nicotinamide adenine dinucleotides (NAD+/NADH) in both chronic and first-episode schizophrenia patients when compared to matched healthy control groups and first-episode bipolar disorder patients [48]. These findings reflect mitochondrial perturbations which are present in schizophrenia at the levels of genes, mRNA, and proteins that negatively impact energy metabolism, redox regulation, antioxidant defense, calcium buffering, and synaptic transmission [18, 19, 48, 55-62]. Furthermore, a transcriptomic study of layer 3 PVIs in the DLPFC of subjects with schizophrenia and unaffected controls indicates that >85% of differentially expressed mitochondrial pathway genes are related to reduced ATP generation and elevated ROS formation [19]. Our findings of increased CypD levels in the DLPFC and within PVIs across layers 2-4 further support the idea that mitochondrial dysfunction and oxidative stress contribute to the etiology of schizophrenia.

Oxidative and other cellular stressors promote the translocation of CypD to the inner mitochondrial membrane and facilitate opening of the mPTP (Figure 5B, [20-22, 28]). CypD has been proposed to interact with several components of the mPTP, including ATP synthase [24, 63], likely binding to the oligomycin sensitivity-conferring subunit of the ATP synthase, promoting a conformational change that leads to the formation of the mPTP [63-65]. Opening of the mPTP rapidly increases the permeability of the inner mitochondrial membrane, collapsing the mitochondrial membrane potential and causing elevated mitochondrial ROS generation and release (Figure 5B, [22, 26, 66, 67]). The mPTP-induced decrease in ATP production can be attributed to the disruption of the proton gradient across the inner mitochondrial membrane, which is essential for ATP synthesis [24, 63, 68]. Consequently, CypD may serve as a unique link between decreased ATP production and increased ROS formation commonly observed in PVIs in schizophrenia, potentially providing a molecular basis for the energy metabolism dysfunction of these neurons in the disorder.

PVIs are particularly susceptible to external stressors during development due to their unique metabolic profile, characterized by a large number of mitochondria and enriched cytochrome-c oxidase [8, 9, 69]. Oxidative stress triggers prolonged CypD-mediated mPTP opening, leading to impaired mitochondrial oxidative phosphorylation and increased ROS formation that may contribute to the observed reduction in PV levels in schizophrenia (Figure 5B). PV serves a critical role in PVIs by regulating intracellular calcium concentrations and fast-spiking, GABAergic inhibitory activity [70, 71]. The mPTP-induced increase in ROS and decrease in ATP production may disrupt cellular energy metabolism, calcium homeostasis, and antioxidant defense mechanisms in PVIs. This disruption could directly or indirectly affect PV expression and function, leading to impaired inhibitory neurotransmission and neuronal synchrony [28, 72, 73]. Our observation of an inverse relationship between CypD and PV in subjects with schizophrenia are consistent with the idea that mitochondrial dysfunction and oxidative stress contribute to reduced PV expression. However, the directionality of these changes is not clear, and alternatively reductions in PV could also lead to increased CypD levels. This notion is supported by the fact that both PV and mitochondria share crucial cellular functions as cytosolic calcium buffers, impact intracellular calcium transients in analogous ways, and jointly maintain calcium homeostasis [74]. Global PV knock-down in mice resulted in a maladaptive increase in mitochondrial volume, likely compensating for the lost calcium buffering capacity of PV, which subsequently contributed to higher oxidative stress levels [75]. Thus, a loss of PV in schizophrenia could trigger a compensatory mitochondrial response, to maintain cellular homeostasis (Figure 5B).

In the DLPFC, neurons in the supragranular layers 2-3 generate gamma oscillations (30-100 Hz) that support cognitive functions like working memory [76, 77]. Functional neuroimaging studies have linked working memory deficits in patients with schizophrenia to diminished gamma oscillatory power in the DLPFC [78-80]. Importantly, working memory deficits and their neural signature are present in the premorbid and prodromal stages of the disease, indicating that DLPFC gamma activity is a strong predictor of functional outcomes in patients who are later diagnosed with schizophrenia [81]. Converging lines of evidence indicate that these prefrontal gamma oscillations depend upon the reciprocal synaptic activity between deep layer 3 PVIs and pyramidal neurons [4, 5, 82-84]. Strong synaptic inhibition from PVIs briefly suppresses pyramidal neuron activity, and once the inhibition decays, the pyramidal neurons fire in synchrony to generate gamma oscillations [83, 84]. The critical role of PVIs is demonstrated by findings that reducing their excitation lowers prefrontal gamma power [85, 86].

However, in schizophrenia pathological alterations also occur in pyramidal cells [82, 87], and transcriptome analyses suggest that mitochondrial impairments might be even more pronounced in pyramidal neurons than PVIs [18, 19]. This may suggest that deficits in pyramidal neurons represent the primary event, followed by compensatory adaptations in PVIs that aim to reduce levels of inhibition. Ultimately, neither mitochondrial deficits or changes in PV may constitute the primary insult, but rather may be a consequence of reduced neuronal demand for energy production due to upstream pathological processes. This notion is supported by postmortem findings which show that layer 3 pyramidal neurons in the DLPFC have fewer dendritic spines and layer 4 PVIs receive less excitatory input, which suggests a primary pathological locus of impaired synaptic processing [85, 87-90]. To better understand the pathology of schizophrenia and develop targeted interventions, it is essential to elucidate the complex interplay and cellular heterogeneity of DLPFC microcircuits, as well as their contribution to gamma oscillations in the disorder.

A general concern for studies using human postmortem tissue is the variability in tissue quality due to perimortem factors such as hypoxia and ischemia, and postmortem factors such as PMI, cause of death, and the conditions under which the tissue was stored and transported [87]. Moreover, the tissue’s integrity may be affected by the process of fixation or embedding, which may distort results across samples [91]. We excluded 12 out of 54 samples (4 control, 8 schizophrenia) because little or no PV signal was detectable in multiple repeats of immunohistochemistry stainings, likely reflecting unspecific factors affecting tissue quality (rather than true differences between diagnostic groups). Another limitation of our study is the inability to control for the potential confounding effects of antipsychotic medications, which are known to cause oxidative stress [92, 93], and thus could influence CypD level. However, previous studies investigating antipsychotic drug effects on PV transcripts [43] and over 600 mitochondrial genes in monkeys exposed to first- and second-generation antipsychotics found minimal differential gene expression [94], suggesting that antipsychotics are unlikely to substantially alter DLPFC functioning through mitochondrial changes. Furthermore, we confirmed that none of the assessed comorbid factors significantly affected PV or CypD protein levels in any our experiments.

In summary, our findings of altered PV and CypD protein levels in the DLPFC suggest that these factors may contribute to redox dysregulation and oxidative stress in cortical PVIs, causing impaired information processing in DLPFC microcircuits in schizophrenia.

## ACKNOWLEDGEMENTS AND DISCLOSURES

This work was supported by grants 1R01 AA028861-01A1 and 1R01 AA028861-01A1 to SK. Human tissue was obtained from the NIH NBB and the Human Brain Collection Core, Intramural Research Program, NIMH (requests #1038 and #1517). We are grateful to the families who gave consent for brain tissue donations. We would also like to thank Dr. Jon Ploski at Penn State College of Medicine for generous donation of western blotting equipment and Dr. Heng Du at the University of Kansas with support in initial studies. The authors declare no conflicts of interest.

